# An integrated workflow for dual-guide CRISPR-Cas9 targeting and amplicon-based genotyping of *Pla2g6* in primary mouse bone marrow-derived macrophages

**DOI:** 10.64898/2026.11.03.763930

**Authors:** Anil K. Challa, Rithvikha Nair, Arumugam Gurusamy, Thi Yen Ly Huynh, Jonathan R. Folmar, Aaliya Ahamed, Sriharsha Chowdarapu, Sasanka Ramanadham

**Affiliations:** Department of Cell, Developmental, and Integrative Biology, University of Alabama at Birmingham, Birmingham, AL 35294, USA; Comprehensive Diabetes Center, University of Alabama at Birmingham, Birmingham, AL 35294, USA

**Keywords:** *Pla2g6*, CRISPR-Cas9, bone marrow-derived macrophages (Mɸs), synthetic guide RNA, heteroduplex mobility assay, targeted amplicon sequencing

## Abstract

CRISPR-Cas9 editing in primary bone marrow-derived macrophages (BMD-Mɸs) requires coordinated assessment of cellular morphology, target modification, and sequence-level editing outcomes. We describe a workflow for dual-guide targeting of murine *Pla2g6* that integrates guide design, ribonucleoprotein nucleofection, PCR-heteroduplex mobility analysis (PCR-HMA), and targeted amplicon sequencing. At 24 h after nucleofection, cells receiving ribonucleoproteins assembled with chemically synthesized single-guide RNAs preserved the elongated, adherent morphology, whereas cultures receiving *in vitro*-transcribed guides contained predominantly poorly adhered, rounded cells and appeared less densely populated; both guide preparations generated indels and deletions, so the difference observed was in cell appearance rather than in target modification. PCR-HMA provided a simple, qualitative screen for sequence heterogeneity and for a putative deletion product, which was subsequently characterized by sequencing. In two independent BMDM samples, one each from C57BL/6 and NOD mice, breakpoint-based analysis identified editing-associated indels in 54.7% and 44.5% of analyzed reads, compared with 0.28% and 0.07% in matched Cas9-only controls. Deletions spanning both target sites represented 33.1% and 26.2% of analyzed reads; none were detected among 4,383 analyzed control reads. The modal spanning deletion was 137 bp, the size predicted from precise joining of the two blunt Cas9 ends. Sequence interpretation depended on verified guide coordinates, deletion-aware analysis settings, and matched controls: scored permissively, the same control reads returned 22-23% apparent editing. Modest-depth Nanopore sequencing recovered the major editing classes, although low-frequency variants and small indels required additional caution. This workflow supports a simple and streamlined screening and sequence-based genotyping of edited BMDM populations, allowing the discernment of the comprehensive mutation spectrum arising from CRISPR-Cas9 gene targeting.

## 1. Introduction

Primary bone marrow-derived macrophages (BMD-M s)are widely used to investigate M biology and gene function. CRISPR-Cas9 ribonucleoprotein (RNP) delivery provides a means of modifying these cells without stable expression of the editing machinery [1]. Successful implementation of the CRISPR-Cas9 gene targeting nevertheless requires attention to cell preparation, guide RNA chemistry and formulation, delivery conditions, and the assays used to characterize the resulting cell population.

While guide RNA and Cas9 can be delivered through plasmid-, viral- and non-viral formats in a variety of cell lines, primary immune cells are sensitive to these formats. *In vitro*-transcribed (IVT) guide RNAs can activate innate immune signaling in susceptible cell types, with effects associated with their 5′-triphosphate groups [3]. The M s are professional sensors of exactly these ligands, so whether a particular guide preparation is suitable for BMDMs requires assessment of the cells after delivery as well as confirmation of target modification.

As CRISPR-Cas9 gene targeting can result in a heterogenous mixture of mutant alleles, there are different methods of performing genotyping populations of cells and resolving the mutation spectrum. PCR-heteroduplex mobility analysis (PCR-HMA) provides an initial qualitative assessment of sequence heterogeneity in an amplified target region. Targeted amplicon sequencing can then identify individual sequence outcomes and estimates their representation among analyzed reads. Interpretation requires explicit definitions of editing classes, appropriate controls, and analysis settings that accommodate the intended deletion. Available analysis tools, including CRISPResso2 [2], facilitate this process, but their inputs and parameters must be matched to the experimental design.

Here, we describe an integrated workflow for targeting *Pla2g6*, which encodes the calcium-independent phospholipase A_2_β (iPLA_2_β), in primary murine BMDMs. The workflow combines dual-guide design, RNP nucleofection, morphological assessment, PCR-HMA screening, and sequence-based genotyping with matched Cas9-only controls. We emphasize practical implementation and the distinction between qualitative screening, read-level editing estimates, and functional gene disruption.

## 2. Materials and methods

### 2.1 Animals and experimental design

All experiments were performed in accordance with the recommendations in the Guide for the Care and Use of Laboratory Animals published by the National Institutes of Health. The protocols used were approved and conducted according to the University of Alabama at Birmingham IACUC (**IACUC Animal Protocol Numbers 22714 and 22535**). Bone marrow was collected from 8-week-old female C57BL/6 or NOD mice. Mice were euthanized by CO_2_ asphyxiation followed by cervical dislocation. Each preparation contributed a dual-guide sample and a matched Cas9-only control.

Three sequencing datasets are reported. Guide chemistry and sequencing platform differ together between them.

**Table 1.**
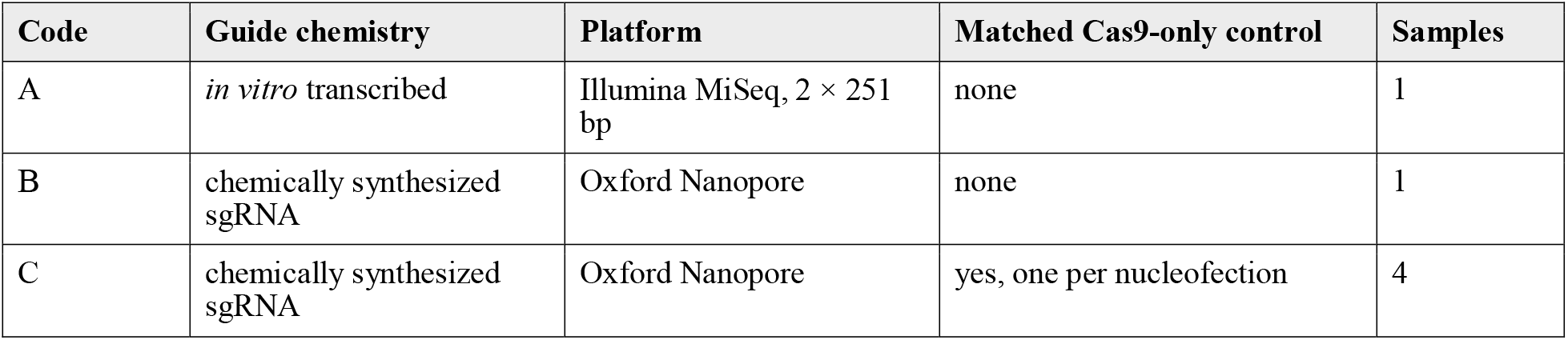
Datasets.

Principal editing estimates were obtained from Dataset C. Datasets A and B were used for exploratory assessment of analysis behavior.

### 2.2 BMDM medium

BMDM medium consisted of 500 mL DMEM/F-12 (1:1) with L-glutamine and 15 mM HEPES (Gibco, 11330-032) supplemented with 50 mL heat-inactivated fetal bovine serum (Phoenix Scientific, PS-300), 5 mL penicillin-streptomycin (Gibco, 15140-122), and 10 µg recombinant murine M-CSF (PeproTech, 315-02). The supplemented formulation yields approximately 555 mL and a final M-CSF concentration of 18 ng/mL; the nominal 20 ng/mL obtained by dividing by the base volume alone is not the working concentration.

### 2.3 Bone marrow isolation and BMDM differentiation

Femurs and tibias were removed, cleared of surrounding muscle and tissue, and placed in 70% ethanol for 1 min. The epiphyses were removed and marrow was flushed with BMDM medium until the bones appeared pale. Solid marrow pieces were dispersed by gentle pipetting, passed through a 70-µm cell strainer, and pelleted at 500 × g for 5 min. Red blood cell lysis was not performed.

Cells from each mouse were resuspended in BMDM medium and divided equally between two T75 flasks containing 10 mL medium per flask, then incubated at 37 °C in a humidified 5% CO_2_ atmosphere for 7 days. Medium was replaced on day 4. By day 7 the cells in each flask were confluent, yielding approximately 8 × 10^6^ BMDMs per flask, or roughly 1.6 × 10^7^ per mouse. On day 7 the medium was removed, the adherent monolayer was washed with warm 1× PBS, and cells were detached with Accutase at 37 °C for 10 min, scraped, and pelleted at 500 × g for 5 min.

Several of these parameters differ from the source protocol for myeloid RNP nucleofection [1].

**Table 2.**
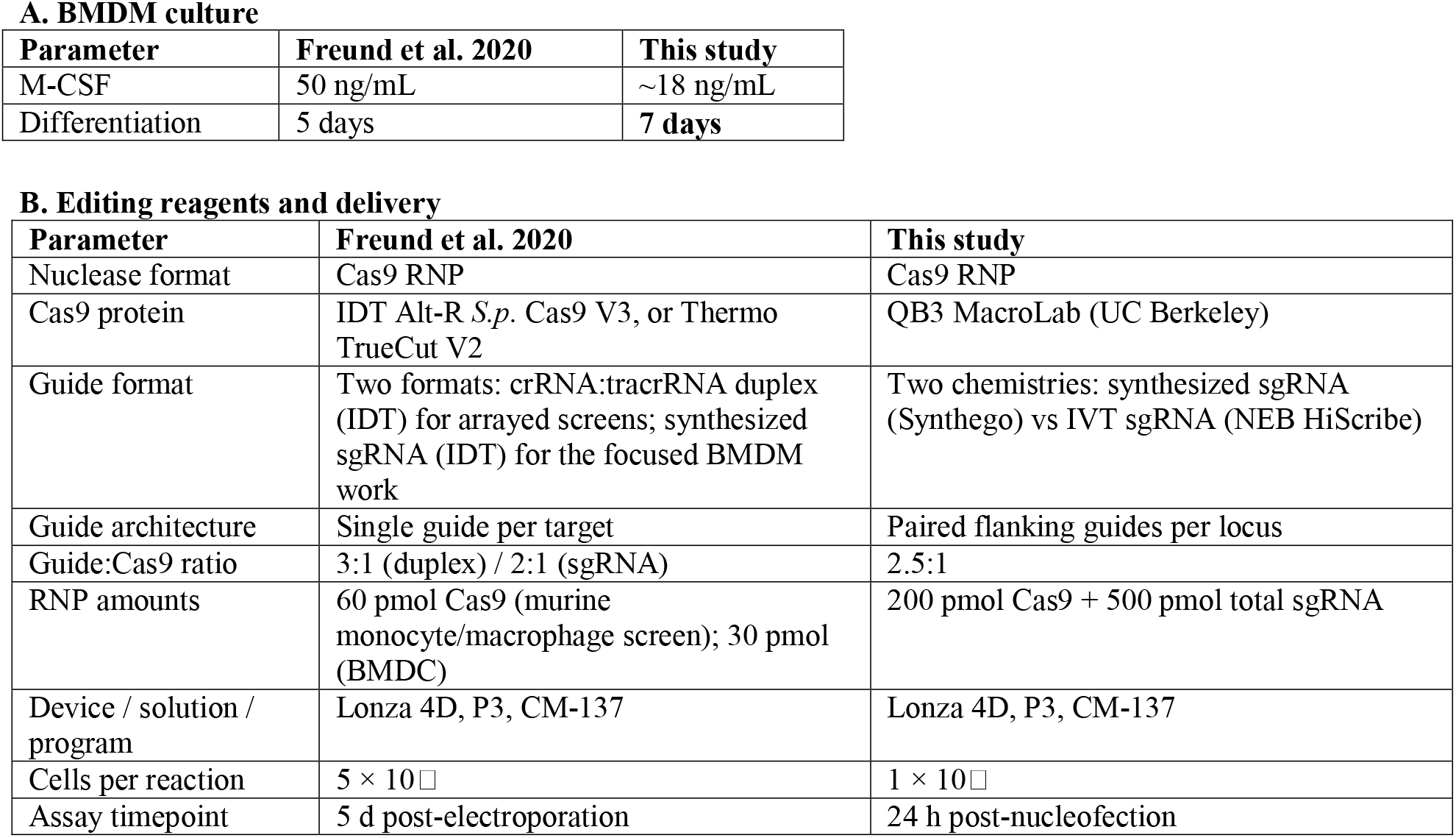
Comparison of the Freund et al. (2020) protocol with the present BMDM gene-targeting workflow.

### 2.4 Target design

Two guides targeted the exon 2 region of *Pla2g6* (ENSMUST00000166977, CCDS27637). Guide 1 had the spacer sequence 5′-CGCTACTGAGGGTGTTGACG-3′ and targets the minus strand and guide 2 had the spacer sequence 5′-GAATGTCTCCAATCGCACCT-3′.

### 2.5 RNP assembly and nucleofection

On day 7 of differentiation, 1 × 10^6^ BMDMs were used per nucleofection with a Lonza 4D-Nucleofector and program CM-137. Each reaction contained 2.5 µL of each chemically synthesized sgRNA at 100 µM (250 pmol per guide, 500 pmol total) and 5.0 µL Cas9 protein at 40 µM (200 pmol), a total guide RNA to Cas9 molar ratio of 2.5:1. Cells were resuspended in P3 Primary Cell Nucleofector Solution supplemented with Supplement 1 before addition of RNPs. Immediately after the pulse, 900 µL of pre-warmed BMDM medium was added to the cuvette and 2-5 × 10^5^ cells were plated per well of a 24-well plate and returned to 37 °C, 5% CO_2_. Cas9-only nucleofections, prepared identically but with Cas9 protein and no sgRNA, were processed in parallel for the Dataset C.

### 2.6 Morphological assessment

Phase-contrast images are acquired 24 h after nucleofection with IVT-guide or synthetic-guide RNPs at 4× and 10× magnification. Cell appearance and apparent density are assessed qualitatively. Viable cell counts were not recorded.

### 2.7 Genomic DNA, PCR and heteroduplex mobility analysis

Cells were lysed 24 h after nucleofection in 50 µl of 50 mM NaOH, neutralized with 2 µl of 1 M Tris-HCl to a final concentration of 40 mM Tris, and 1 µL of the crude lysate was used directly as PCR template. The target region was amplified using Mm_*Pla2g6*-Ex2-gen-F2 (5′-TTCTTCCAGAGCAGAGGGGCAGG-3′) and Mm_*Pla2g6*-Ex2-gen-R2 (5′-AACGAACTGAGGGGTGCACGTC-3′) with NEB 2× Taq Polymerase Master Mix, yielding a 298 bp reference product.

PCR products were analyzed directly by heteroduplex mobility assay on 6% TBE-polyacrylamide gels. Heteroduplex species, formed when wild-type and indel-containing strands reanneal during the final cycles of PCR, migrate more slowly than the homoduplex product and indicate a mixed allele population; the shorter programmed excision product resolves as a discrete band.

### 2.8 Targeted amplicon sequencing

Dataset A was generated by MR DNA (Shallowater, TX) using a tag-encoded amplicon approach on Illumina MiSeq with paired-end 2 × 251 bp reads. Synthetic-guide amplicons (Datasets B and C) were sequenced by Plasmidsaurus using their amplicon Edit-efficiency service for small edits. Dataset C comprised four samples from two nucleofections.

### 2.9 Read accounting and exclusion rules

Reads were processed from the vendor FASTQ without trimming or error correction. Reads longer than 700 nt were discarded as concatemers or chimeras, as were reads with a mean Phred score below 12. Reads were accepted in either orientation and reverse-complemented into reference orientation using the left anchor as probe. Reads in which an anchor failed to match were recorded as a separate non-scorable class and were never assigned to an allele; this dropout is non-random and is enriched for indels, so assignment would bias the estimate. Substitutions were never counted as editing.

Illumina reads were verified by checksum on download and re-paired with seqkit after an initial download produced truncated and mispaired files.

### 2.10 Allele calling and editing quantification

Two complementary approaches were used for allele calling and editing quantification. A breakpoint-based analysis assigned each indel event to either target site, or to a deletion spanning both sites, according to its reference interval, and scored a read as edited only when a breakpoint fell within ±10 bp of a predicted cut site. A separate anchor-based analysis placed 20 bp reference anchors 5 bp outside each cut site and classified the net change in the distance between them; for allele calling in Dataset C a three-anchor geometry was used, with 24 bp anchors at positions 21-44, 130-153 and 240-263, and a read was called as carrying the programmed excision when the middle anchor was absent and the outer span had shortened by at least 50 bp.

The excision test was applied to every allele independently of the per-site classification, with 2 bp of end-processing tolerance, because a deletion beginning one position inside the cut is the same programmed event with additional resection. Reading frame was assigned from the net coding-length change. Reads were subsampled without replacement at increasing depths, with 40 replicates per depth, to describe variability in the recovered editing classes. Analyses performed under different criteria are reported separately and are not combined.

As an independent check on implementation, the Dataset B was additionally analyzed with a hybrid method that used affine-gap global alignment for large deletions and paired anchors for per-site indel sizing.

### 2.11 Background estimation

Background was estimated in two ways. The matched Cas9-only control was scored at the actual cut sites under identical criteria and provides the correct factor for comparison. In the absence of such a control, four no-cut windows within the same amplicon, at positions 30, 140, 170 and 265, were scored with identical anchor geometry. Both estimates were computed on the same control reads so that the two could be compared directly.

## 3. Results

### 3.1 Implementation of the workflow and expected target modification

The workflow integrates BMDM differentiation, dual-guide RNP nucleofection, morphological assessment, PCR-HMA, and targeted amplicon sequencing, with sequence interpretation resting on verified guide coordinates, explicit variant classification, and comparison with matched controls (**Fig. 1**). The guide pair was positioned to generate a readily distinguishable deletion product within a single genotyping amplicon, so that a gel-based screen and a sequencing-based measurement interrogate the same event.

**Figure 1.**
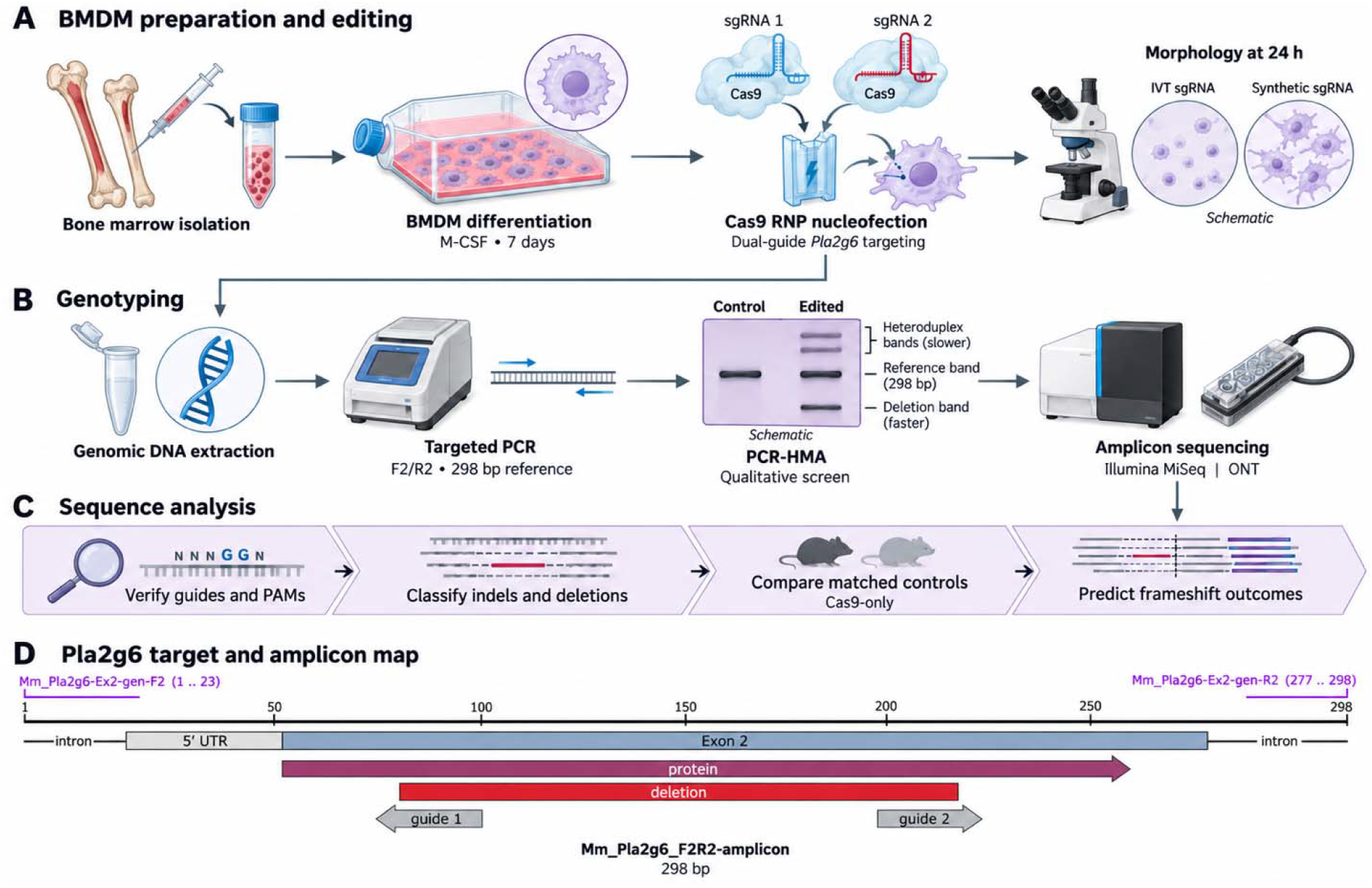
Workflow for dual-guide *Pla2g6* targeting and genotyping in primary BMDMs. **(A)** BMDM preparation and editing. Bone marrow is isolated from femurs and tibias, differentiated for 7 days in M-CSF, and nucleofected on day 7 with Cas9 ribonucleoproteins carrying two sgRNAs against *Pla2g6*; cultures are imaged at 24 h to compare *in vitro*-transcribed with chemically synthesized sgRNA. **(B)** Genotyping. Genomic DNA is amplified with the F2/R2 primer pair to give a 298 bp reference product, screened by PCR-HMA for slower-moving heteroduplex species and for the shorter deletion product, and then sequenced by Illumina or Oxford Nanopore amplicon sequencing. **(C)** Sequence analysis. Guides and PAMs are verified against the reference before quantification windows are defined; indels and spanning deletions are classified explicitly; matched Cas9-only samples provide the background against which editing is scored; and reading-frame outcomes are predicted from coding-length change. Gel and morphology elements in panels A and B are schematic. **(D)** Annotated amplicon map showing the 298 bp coordinate frame, the ATG at 51-53, the cut sites at 73|74 and 210|211, and the 137 bp excision.

### 3.2 Synthetic guides are associated with preserved adherent morphology of M s

At 24 h after nucleofection, cultures receiving IVT guides appeared less densely populated and contained predominantly rounded, refractile cells, whereas cultures receiving chemically synthesized sgRNAs retained the elongated and adherent cells with cellular extensions in the displayed fields (**Fig. 2**). The rounded morphology was persistent and had an impact on cell growth and proliferation over subsequent days. Wienert et al., [3] show that the IVT guides used for RNP-based targeting trigger a potent innate immune response. They further show that the removal of the 5’-triphosphate from gRNAs ameliorates inflammatory signaling and prevents the loss of viability associated with genome editing in hematopoietic stem cells.

**Figure 2.**
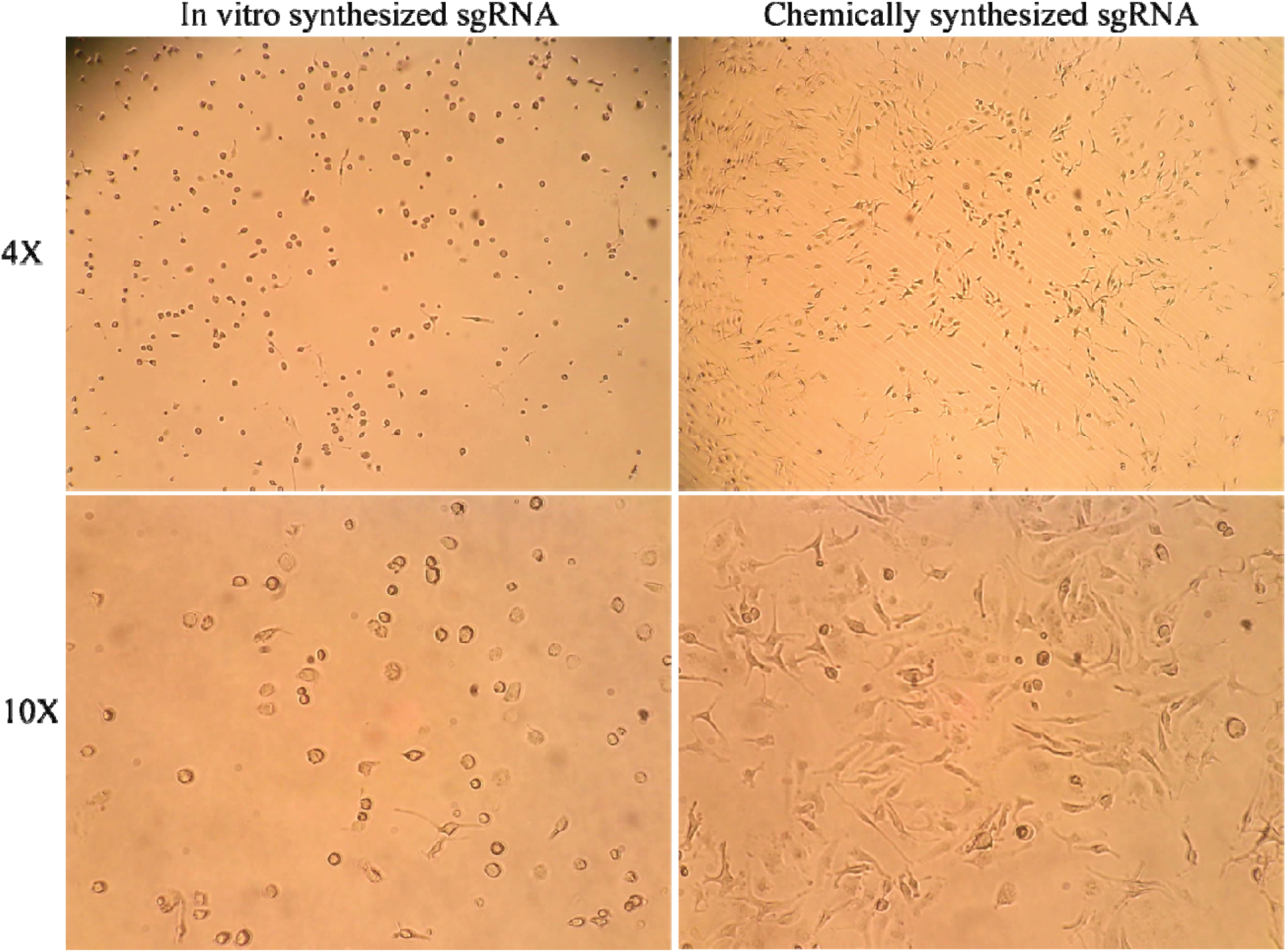
BMDM morphology 24 h after nucleofection with IVT or chemically synthesized guide RNAs. Phase-contrast images at 4× and 10× magnification. IVT-guide cultures contain predominantly rounded cells and appear less densely populated; synthetic-guide cultures retain more spread, adherent cells.

Independent of the effect on cell morphology, both preparations generated indels and deletions at the target sites in BMDMs, so the difference between them is in cell morphology rather than in whether the RNP cut the target sites effectively.

To ensure healthy growth and proliferation of BMDMs nucleofected with the CRISPR-Cas9 RNP complex, the use of chemically synthetic guides, in contrast to *in vitro* synthesized guides, is an essential factor.

### 3.3 PCR-HMA identified samples for sequence-based characterization

PCR-HMA revealed slower-moving products above the reference-sized band in the dual-guide sample, together with a shorter band at approximately the size expected for the programmed deletion product (**Fig. 3**). These bands correspond to heteroduplexes formed by complementary strands of wildtype and indel mutant sequences. This observation has been reported previously [4,5,6]. The corresponding heteroduplex features were not evident in the displayed Cas9-only lane. These findings gave credence and supported progression to sequence-based analysis. PCR-HMA is an easy and quick qualitative screening step to help decide to progress to sequence-based analysis.

**Figure 3.**
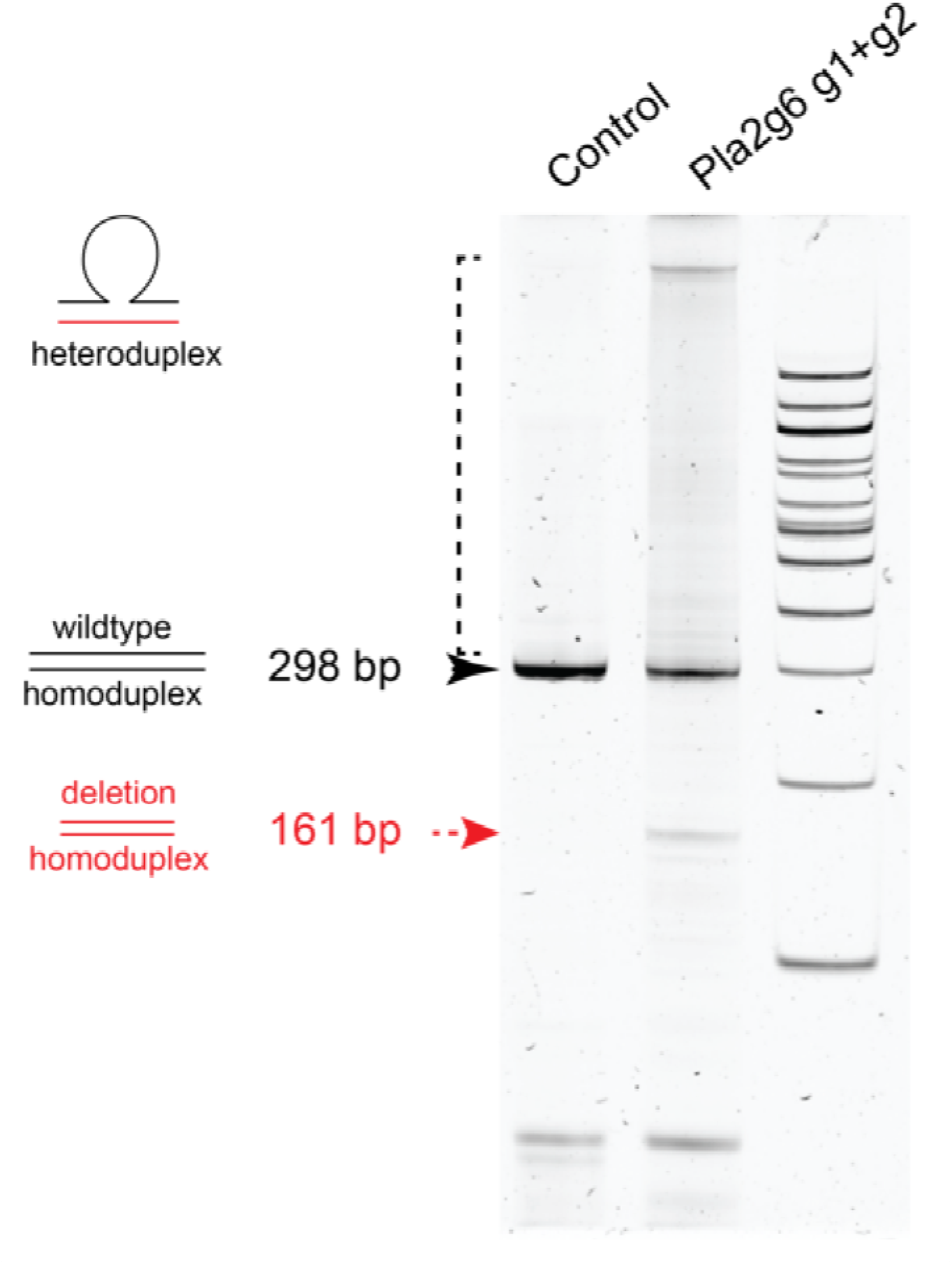
PCR-HMA screening of a dual-guide *Pla2g6* sample. Slower-moving products are visible above the 298 bp reference product in the dual-guide lane, together with a shorter product, 161 bp, migrating near the size expected for the 137 bp deletion.

### 3.4 Sequencing identified editing-associated indels and deletions spanning both targets

As the sequencing was performed on the 298 bp genotyping amplicon, all coordinates refer to this reference sequence. The initiating ATG occupies positions 51-53 and the coding sequence within the amplicon spans positions 51-260.

#### Streptococcus pyogenes

Cas9 protein cleaves between positions 17 and 18 of the spacer, 3 bp from the PAM. Therefore, the blunt cuts are between *Pla2g6* amplicon positions 73 and 74 for guide 1 and between 210 and 211 for guide 2. Precise joining of the two blunt ends removes positions 74-210, a 137 bp deletion, and yields a 161 bp product. Positions 74 and 211 are both C, so the junction carries 1 bp of microhomology and the identical sequence can be written as a deletion of 74-210 or of 75-211; the left-aligned form is used throughout because it coincides with the cut sites.

Using this as a reference framework, in the Dataset C, breakpoint-based analysis identified editing-associated indels in 54.7% of analyzed reads from the C57BL/6 samples and 44.5% from the NOD sample. Corresponding Cas9-only values were 0.28% and 0.07%. Deletions spanning both target sites accounted for 33.1% and 26.2% of analyzed reads, with no such events detected among 4,383 analyzed control reads (**Fig. 4A**; **Table 3**).

**Table 3.** Editing in two BMDM preparations with matched Cas9-only controls (Dataset C). Percentages are of analyzed reads. Frameshift and in-frame classes are predictions from coding-length change, not measurements of protein loss.

| **Sample** | **Analyzed reads** | **Total edited** | **Spanning deletion** | **Predicted frameshift** | **Predicted in-frame** |
|---|---|---|---|---|---|
| Cas9-only, Nuc #5 (C57BL/6) | 1,417 | 0.28% | 0.00% | 0.28% | 0% |
| *Pla2g6* RNP, Nuc #5 (C57BL/6) | 1,752 | 54.7% | 33.1% | 34.8% | 20.0% |
| Cas9-only, Nuc #6 (NOD) | 2,966 | 0.07% | 0.00% | 0.07% | 0% |
| *Pla2g6* RNP, Nuc #6 (NOD) | 2,380 | 44.5% | 26.2% | 31.5% | 12.9% |

**Figure 4.**
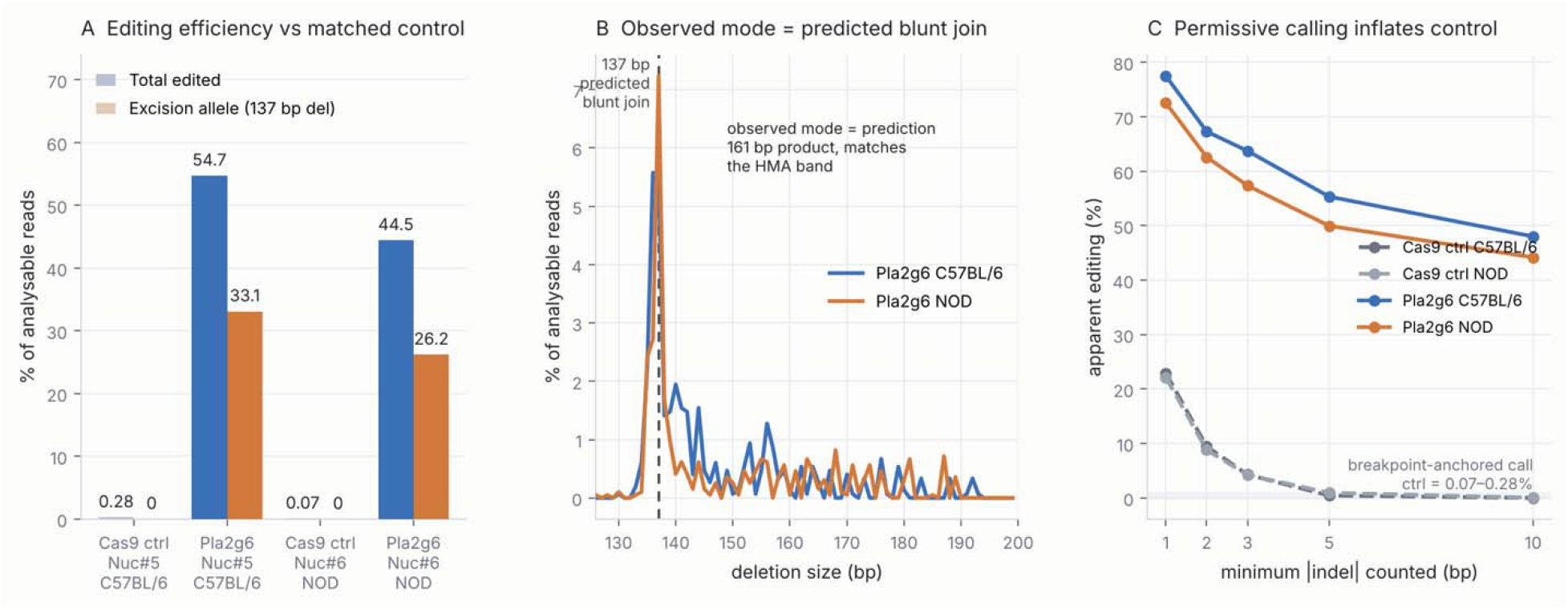
Breakpoint-based editing analysis in two BMDM samples with matched controls. **(A)** Editing-associated indels and deletions spanning both target sites, in edited samples and matched Cas9-only controls, as a percentage of analyzed reads. **(B)** Length distribution of deletions spanning both sites; the dashed line marks the 137 bp product predicted from precise blunt joining. **(C)** Apparent editing as a function of the minimum indel size counted, shown separately from the breakpoint-based classification.

The expected 137 bp deletion was the modal spanning-deletion length in both samples (**Fig. 4B**). A 161 bp contig was assembled independently from each edited sample; the two are identical to one another at every position and are reconstructed exactly as reference positions 1-73 joined to 211-298, matching the product predicted from the two RNP cut sites and the size of the discrete band seen by HMA. An independent anchor-based recount of the raw reads returned spanning-deletion frequencies of 36.7% and 31.0% of callable reads, within the same range on a different denominator.

As multiple spanning-deletion lengths were observed, the overall spanning-deletion frequency should be distinguished from the frequency of the exact predicted junction. In the Dataset B, where junctions were enumerated directly, the predicted blunt join was the single most abundant spanning-deletion allele and the most frequent junctions carried no microhomology, consistent with blunt non-homologous end joining rather than microhomology-mediated end joining.

The Cas9-only consensus sequences from the C57BL/6 and NOD samples are identical at all 298 amplicon positions, including both protospacers, both PAMs and both primer sites. Strain cannot therefore confound these measurements through target-site or primer-site sequence.

### 3.5 Guide coordinates and matched controls determined the interpretation

Analysis configuration affected both the placement of quantification windows and the classification of sequence outcomes. In one configuration, the plus-strand representation of the reverse-strand guide placed the window 14 bp from the expected cleavage position, and the analysis reported a result rather than an error. Separately, a vendor analysis that inferred the spacer from the data placed guide 2 one nucleotide from the experimentally supplied sequence; both registers are legitimate NGG sites, but they predict products differing by one base. Verification of the actual spacer sequences, strand orientation and PAMs resolved both discrepancies, and the assembled 161 bp product is consistent with the verified coordinates.

Matched controls revealed substantial background under permissive calling criteria. Counting any net length change across the cut-site interval, Cas9-only control reads returned 22.9% and 22.2% apparent editing. Requiring indels of at least 3 bp reduced this to 4.3% and 4.2%, at least 5 bp to 0.45% and 0.95%, and at least 10 bp to 0.0% and 0.04%. The corresponding dual-guide samples remained above 44% at every threshold (**Fig. 4C**; **Table 4**). Permissive calling is therefore not the conservative choice; it generates a signal, and the size of the effect is not visible without a control.

**Table 4.** Apparent editing as a function of the minimum indel size counted (Dataset C, anchor-based criterion). Percentages are of scorable reads.

| **Sample** | **≥1 bp** | **≥2 bp** | **≥3 bp** | **≥5 bp** | **≥10 bp** |
|---|---|---|---|---|---|
| Cas9-only, C57BL/6 | 22.9% | 9.5% | 4.3% | 0.45% | 0.0% |
| Cas9-only, NOD | 22.2% | 8.8% | 4.2% | 0.95% | 0.04% |
| *Pla2g6* RNP, C57BL/6 | 77.5% | 67.3% | 63.7% | 55.3% | 48.0% |
| *Pla2g6* RNP, NOD | 72.6% | 62.6% | 57.4% | 50.0% | 44.2% |

Background noise also varied with position in the amplicon. In the C57BL/6 control, apparent indel frequencies at four no-cut windows chosen ranged from 1.20% at position 265 to 9.53% at position 30, an eightfold spread, with a mean of 4.51%; the values measured at the actual cut sites 1 and 2 in the same reads were 1.48% and 2.29%. In the NOD control the no-cut mean was 3.59% against cut-site values of 1.12% and 1.32% (**Fig. 5A**). The within-amplicon surrogate thus overestimated cut-site background two-to threefold and varied several-fold with window placement. It is a conservative bound rather than a measurement, and these observations favor measuring background at the actual target sites in matched samples over assuming that other positions carry equivalent error characteristics.

**Figure 5.**
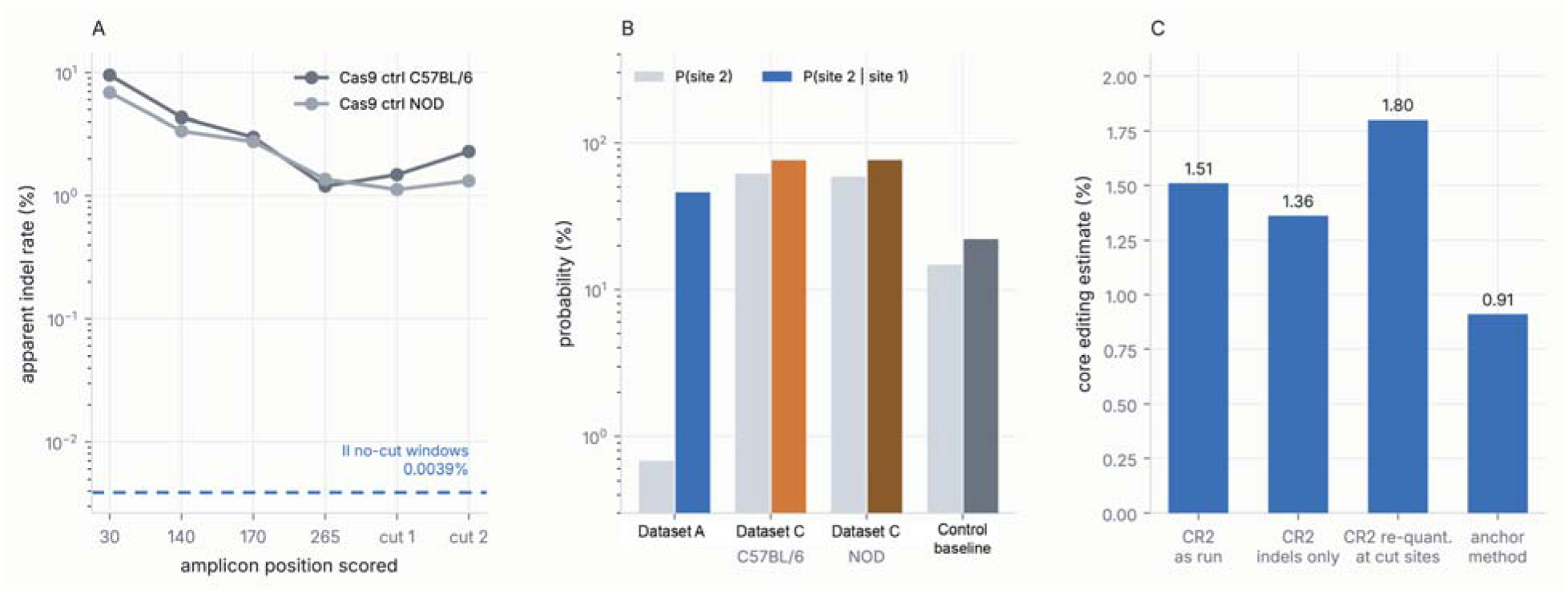
Effects of background position and analysis criteria. **(A)** Apparent indel frequencies at four no-cut windows and at the two target sites, measured in matched Cas9-only controls. **(B)** Marginal and condition frequencies of detectable editing-associated outcomes. **(C)** Core editing estimates obtained from a single Illumina dataset under four defensible analysis configurations.

### 3.6 Editing classes and predicted frameshift outcomes are distinct quantities

Anchor-based analysis separated single-site changes, independent changes at both sites, and deletions spanning the target interval (**Fig. 6A**). Per-site distributions included a prominent 3 bp deletion at site 1 and small deletions and insertions at site 2 (**Fig. 6B,C**). Similar prominent indel sizes were observed in the two samples, although the design does not establish strain-independent repair distributions.

**Figure 6.**
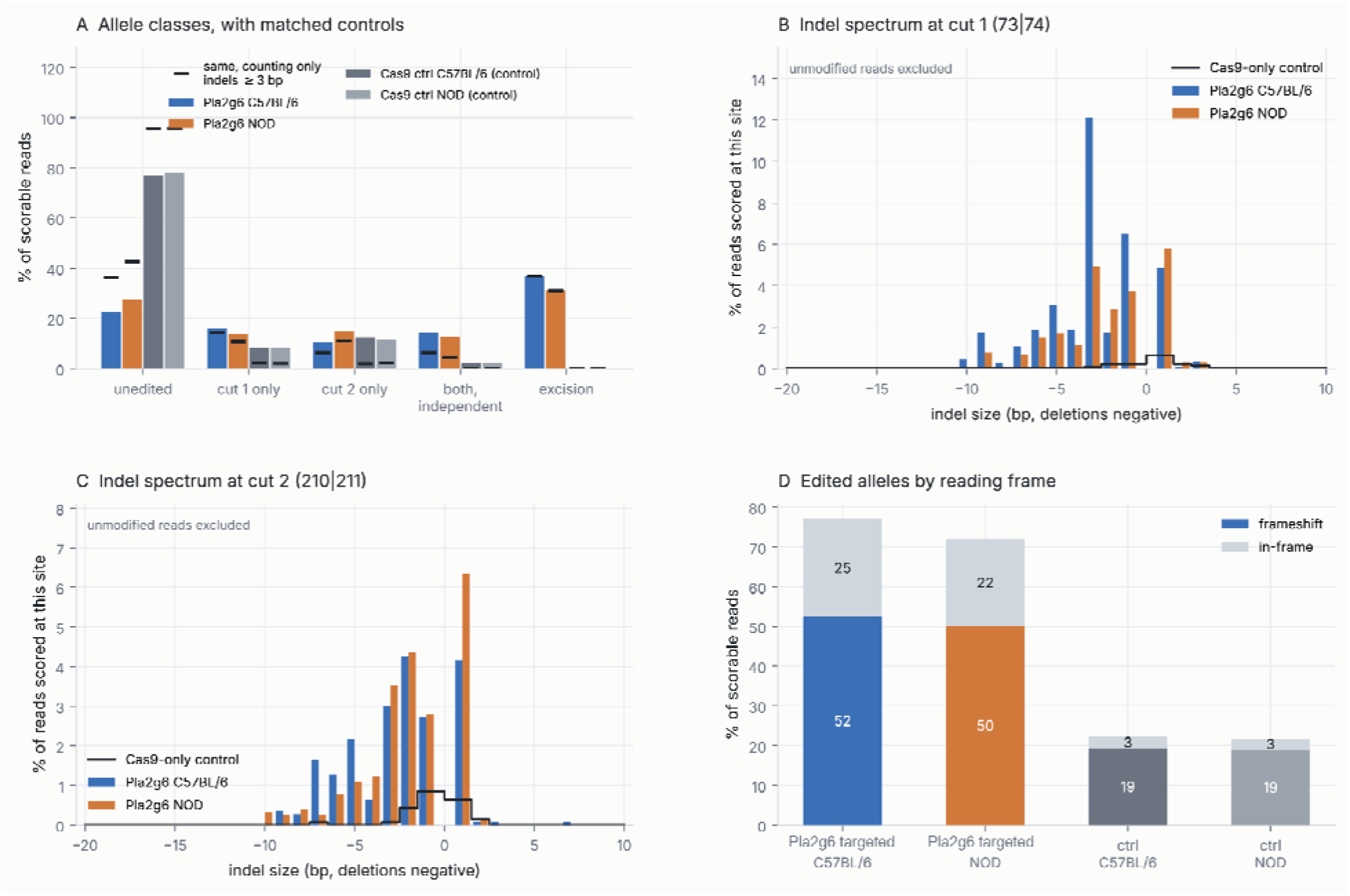
Anchor-based classification of sequence outcomes in Dataset C. **(A)** Read classes under permissive and minimum 3 bp indel criteria, with matched Cas9-only controls. **(B,C)** Per-site indel-length distributions at cut 1 (73|74) and cut 2 (210|211), unmodified reads excluded, with the Cas9-only control spectrum overlaid. **(D)** Reading-frame classes under the anchor-based criterion, including matched controls. The estimates in (D) obtained under a different operational definition from the breakpoint-based frameshift frequencies reported in Section 3.6 and are not interchangeable with them.

Breakpoint-based reading-frame analysis yielded predicted frameshift frequencies of 34.8% and 31.5%, against total editing estimates of 54.7% and 44.5%. Total editing exceeded predicted frameshift frequency by 19.9 and 13.0 percentage points, the difference being in-frame alleles at 20.0% and 12.9%. These results distinguish target modification from predicted coding disruption. They do not quantify functional-null alleles, nor the proportion of cells carrying loss-of-function alleles on both chromosomes, and whether the in-frame class removes catalytically required residues was not determined.

### 3.7 Modest sequencing depth recovered the common outcome classes

Nanopore sequencing recovered the major editing classes in these efficiently edited samples from a few thousand raw reads per sample. Repeated subsampling produced similar mean class proportions, with greater variability at lower depths, and the class proportions were stable from approximately 1,000 reads upward (**Fig. 7A**). The number of distinct recovered indel lengths continued to increase with depth (**Fig. 7B**), indicating that the observed length distribution was incompletely sampled even at full depth.

**Figure 7.**
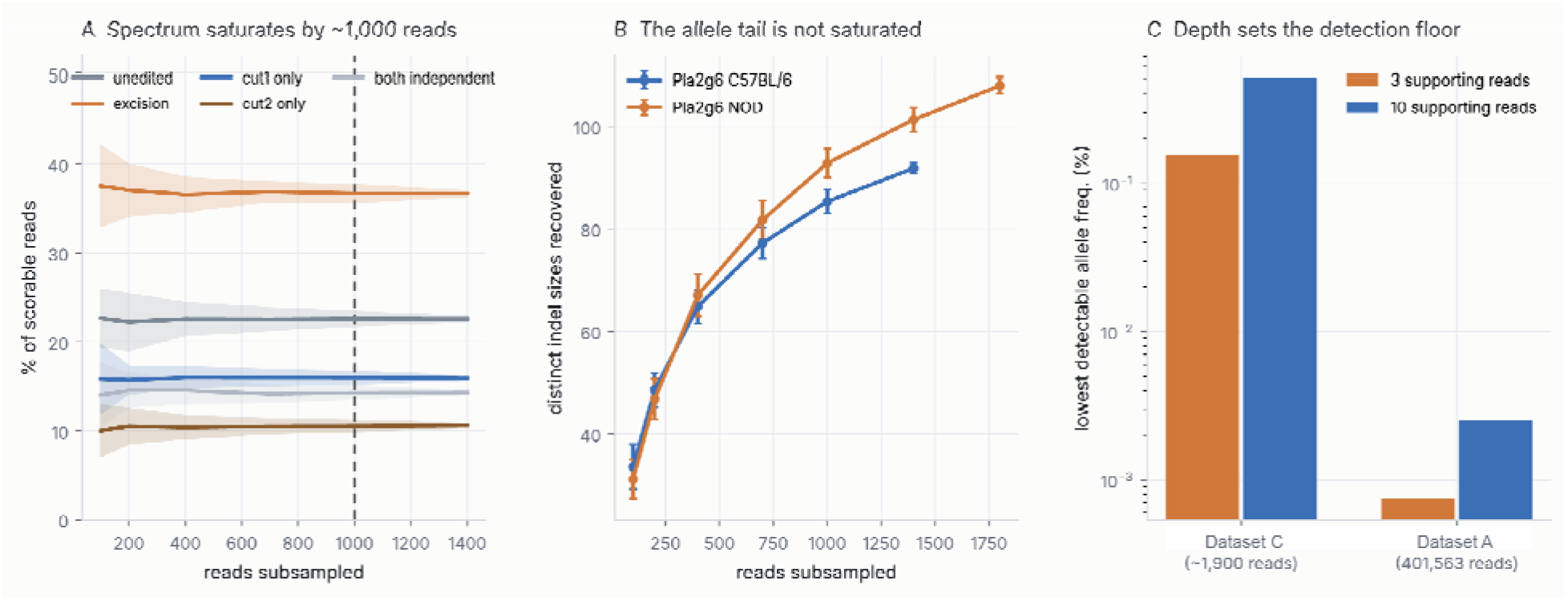
Effect of sequencing depth, Dataset C. **(A)** Mean editing-class proportions with ±1 SD across 40 subsamples per depth. **(B)** Number of distinct indel lengths recovered as a function of depth. **(C)** Lowest detectable allele frequency corresponding to a specified number of supporting reads. Subsampling variability describes sampling behavior at the achieved depth; it does not establish biological reproducibility or a validated limit of detection.

These results support practical characterization of common editing outcomes at the achieved depths. They do not establish a universal minimum depth, nor a validated limit of detection for rare variants. Frequencies calculated from a specified expected read count describe nominal sampling support only, and must be interpreted together with background, error classification, and the detection probability required for the question at hand.

## 4. Discussion

This study describes an integrated workflow for CRISPR-Cas9 targeting and amplicon-based genotyping of *Pla2g6* in primary BMDMs. The workflow connects experimental design to cellular assessment, inexpensive gel screening, and sequence-level characterization. Its practical value lies in defining what each stage establishes and when additional analysis is required.

Synthetic sgRNAs were associated with better preservation of elongated, adherent morphology than IVT guides in the displayed comparison. Previous work demonstrated that IVT guide RNAs can stimulate RIG-I-dependent responses in susceptible cells [3], and M s are professional sensors of the 5′-triphosphate and double-stranded RNA species that IVT preparations retain, which offers a plausible explanation for the observed difference.

PCR-HMA provided a simple, easy and reliable first-stage screen. Its role is complementary to sequencing: altered mobility identifies sequence heterogeneity, whereas sequence analysis identifies the underlying variants and their junctions. A shorter band at the expected size supports a candidate deletion but does not establish its identity conclusively. This staged approach can prioritize samples for sequencing without treating band intensity as an editing percentage.

One observation deserves emphasis because it bears on how amplicon libraries are read. HMA is performed on unpurified PCR product and showed a comparable deletion band in both the IVT and synthetic experiments, whereas the Illumina library from the IVT experiment reported the spanning deletion at a small fraction of a percent. The two mechanisms underestimate the short-read value and both act in the same direction: default global-alignment gap penalties divert the 137 bp deletion out of the output, and size-selective bead cleanup during library preparation depletes a 161 bp product that gel-based and long-read methods, which interrogate the raw PCR product, both retain. The Illumina spanning-deletion value is therefore only a safe upper limit, and should not be quoted as an editing efficiency or compared directly with the Nanopore figures.

Targeted amplicon sequencing provides a practical route to detailed genotyping through external services, including for laboratories without dedicated sequencing instruments. In the present study, Nanopore reads were sufficient to recover the major editing classes and the predicted dual-cut junction. Previous work has also evaluated Nanopore-based CRISPR indel profiling [7]. Accessibility depends on sample requirements, usable read recovery, and the availability of a reproducible analysis workflow.

It is important to note that the analysis should be tailored to the designed outcome. Guide sequences and PAM orientation must be verified before quantification windows are selected, and the analysis must accommodate a deletion of the expected length. Matched controls are particularly useful for separating low-frequency and small-indel signals from assay background. In our hands, the common alternative of estimating background from other positions in the same amplicon overestimated it two-to threefold and varied eightfold with window placement. Therefore, multiple analysis methods can reveal sensitivity to assumptions, but outputs obtained under different denominators or definitions should not be combined into a single uncertainty interval.

Editing frequency and predicted frameshift frequency address different aspects of the edited population and should each be reported with their definition and denominator. Functional disruption requires additional evidence appropriate to the biological question, such as assessment of protein expression or activity. Bulk amplicon sequencing does not assign alleles to individual cells and therefore cannot establish the proportion of cells carrying loss-of-function alleles on both chromosomes.

The workflow was demonstrated at one locus in two independent samples from different mouse backgrounds, with one preparation per background. The morphology comparison was qualitative. The three sequencing datasets differ in several experimental and analytical factors simultaneously, so none of the comparisons between them is controlled. PCR amplification, library processing and read filtering may alter observed proportions relative to genomic allele frequencies. It is pertinent to note that shorter deletion products are expected to amplify preferentially during PCR, making spanning-deletion frequencies an upper bound on the true allele frequency in the cell population. These limitations restrict broader generalizations, but do not discount or negate the feasibility of integrating morphology, PCR-HMA screening and targeted sequencing into BMDM editing experiments using the CRISPR-Cas9 system.

## 5. Conclusion

Dual-guide *Pla2g6* targeting in primary BMDMs can be evaluated through a staged workflow comprising morphological assessment, PCR-HMA screening, and targeted amplicon sequencing. Synthetic guides were associated with preservation of elongated, adherent morphology under the examined conditions, while both guide chemistries produced indels and deletions. PCR-HMA identified candidate edited samples, and sequencing characterized their editing-associated indels and the predicted dual-cut junction. Verified guide coordinates, matched Cas9-only controls, deletion-aware alignment settings, and explicit read-level definitions are each necessary for reliable interpretation. The failure of any one of these produces a plausible number rather than an error.

## Declarations

### Author contributions

Contributions are described using the CRediT taxonomy. **Anil K. Challa:** Conceptualization, Methodology, Software, Formal analysis, Investigation, Data curation, Visualization, Supervision, Writing - original draft, Writing - review & editing. **Rithvikha Nair:** Investigation, Validation, Writing - review & editing. **Arumugam Gurusamy:** Investigation, Writing - review & editing. **Thi Yen Ly Huynh:** Validation, Investigation. **Jonathan R. Folmar:** Validation, Writing - original draft, Writing - review & editing. **Aaliya Ahamed:** Investigation, Writing - review & editing. **Sriharsha Chowdarapu:** Investigation, Writing - review & editing. **Sasanka Ramanadham:** Conceptualization, Supervision, Project administration, Funding acquisition, Resources, Writing - review & editing. All authors read and approved the final manuscript.

### Use of Artificial Intelligence (AI)

Generative AI was used in the preparation of this work and is disclosed here in full. Claude (Anthropic) Opus 5 was used for three specific purposes: reanalysis of previously generated sequencing data; generation of the read-classification, background-estimation and figure-generation code deposited with this manuscript; and editing of the manuscript text.

AI tools were not used to generate, acquire or alter any primary data. All sequencing reads, gel images and micrographs reported here were produced experimentally by the authors. No artificial intelligence tool is credited with authorship, in accordance with the position that authorship entails accountability that such tools cannot assume.

## Funding

This work was supported by the following grants to S.R. - Breakthrough T1D: INO-2023-1344-A-N; SRA-2023-1362-S-B; NIH/NIAID: NIH/NIAID 1R21AI169214-01; Beatson Foundation #2023-20; UAB Diabetes Research Center P30 DK079626

## Acknowledgments

The authors thank Ying Gai Tusing for animal care and laboratory support; Xiaoyong Lei and members of the Comprehensive Diabetes Center at the University of Alabama at Birmingham; and Dr. Rosa Serra for help with nucleofection.

## Competing interests

The authors declare no competing interests.

## Ethics approval

All animal procedures were approved by the Institutional Animal Care and Use Committee of the University of Alabama at Birmingham (**IACUC protocol number**) and were performed in accordance with the Guide for the Care and Use of Laboratory Animals published by the National Institutes of Health.

## Data availability

Raw sequencing reads have been deposited in the NCBI Sequence Read Archive under BioProject accession **PRJNA1554390.**

## Code availability

All analysis code, reference files, the target configuration and the exact commands used to generate the reported values are available at https://github.com/anilkchalla/pla2g6-bmdm-crispr-analysis, and released under the MIT license.

## Notes

https://www.ncbi.nlm.nih.gov/bioproject/PRJNA1554390

https://github.com/anilkchalla/pla2g6-bmdm-crispr-analysis

### Competing Interest Statement

The authors have declared no competing interest.

### Summary of Updates

Order of authors is revised to reflect the list in the manuscript.

## References

1. Freund EC, Lock JY, Oh J, et al. Efficient gene knockout in primary human and murine myeloid cells by non-viral delivery of CRISPR-Cas9. J Exp Med. 2020;217(7):e20191692. 10.1084/jem.20191692

2. Clement K, Rees H, Canver MC, et al. CRISPResso2 provides accurate and rapid genome editing sequence analysis. Nat Biotechnol. 2019;37:224–226. 10.1038/s41587-019-0032-3

3. Wienert B, Shin J, Zelin E, Pestal K, Corn JE. *In vitro*-transcribed guide RNAs trigger an innate immune response via the RIG-I pathway. PLoS Biol. 2018;16(7):e2005840. 10.1371/journal.pbio.2005840

4. Barlow, K. L., J. Green, and J. P. Clewley. “Viral genome characterisation by the heteroduplex mobility and heteroduplex tracking assays.” Reviews in medical virology 10.5 (2000): 321–335.

5. Challa, A. K., Boitet, E. R., Turner, A. N., Johnson, L. W., Kennedy, D., Downs, E. R., … & Kesterson, R. A. (2016). Novel hypomorphic alleles of the mouse tyrosinase gene induced by CRISPR-Cas9 nucleases cause non-albino pigmentation phenotypes. PLoS One, 11(5), e0155812.

6. Foster, S. D., Glover, S. R., Turner, A. N., Chatti, K., & Challa, A. K. (2019). A mixing heteroduplex mobility assay (mHMA) to genotype homozygous mutants with small indels generated by CRISPR-Cas9 nucleases. MethodsX, 6, 1–5.

7. McFarlane GR, Polanco JVC, Bogema D. CRISPR-Cas guide RNA indel analysis using CRISPResso2 with Nanopore sequencing data. BMC Res Notes. 2024;17:205. 10.1186/s13104-024-06861-1

